# The impact of colonization history on the composition of ecological systems

**DOI:** 10.1101/2020.02.26.965715

**Authors:** Nannan Zhao, Serguei Saavedra, Yang-Yu Liu

## Abstract

Observational studies of ecological systems have shown that different species compositions can arise from distinct species arrival orders during community assembly—also known as colonization history. However, it is still unclear under which conditions colonization history will dominate community composition. Yet, this is important in order to understand and anticipate the impact of species arrivals on the biodiversity that we observe in both nature and experiments. To address this fundamental question, here we develop a testable theory linking colonization history and community composition. First, we prove two general theorems to determine whether the composition of a community will depend on its colonization history. For communities governed by Lotka-Volterra dynamics, we further simplify the two theorems into a corollary that is easy for numerical test. Second, we show, via extensive numerical simulations, that the probability that community composition is dominated by colonization history increases monotonically with community size and species connectivity. Third, we show that this probability significantly increases with the variation of intrinsic growth rates across species. These results reveal that community composition is a probabilistic process mediated by ecological dynamics via the interspecific variation and the size of regional pools.

---

Ecological communities are formed by co-occurring and interacting species in a given place and time [1–3]. It has been shown that within these communities, the specific composition of species is a function of several ecological, evolutionary, and stochastic processes [3–6]. Importantly, one of the main factors affecting community composition is the order of species arrival—also known as colonization history [7–11]. That is, colonization history can introduce priority effects, where the persistence of species depends on the order at which they join a given community. However, it remains unclear under which conditions can colonization history have the strongest impact on the assembly of ecological communities [12–15]. For example, are there any conditions under which colonization history will completely dominate community composition? Does the type of interspecific interactions affect the probability that community composition depends on colonization history? How do the intrinsic properties of species affect the impact of colonization history on community composition? In the face of an accelerating rate of species turnover, answering these questions is important in order to understand and anticipate key biodiversity changes in ecological communities.

To answer the questions above, previous work has established different null models and compared the composition of observed communities against what would be expected by chance alone under different colonization histories [16–19]. These null models have ranged from mechanistic processes to naive random assemblages. For instance, mechanistically, it has been shown that the trophic level [20], environmental feedback [21], and the rate of nutrient supply [22] can determine the extent to which community composition depends on colonization history. Theoretically, it has also been shown that community composition strongly depends on colonization history [23–26]. Nevertheless, these approaches do not allow us to find general conditions under which colonization history can have the highest (or lowest) chance to affect community composition. Moreover, the complexity of factors affecting community assembly has undercut our ability to anticipate whether a given regional pool of species can be more susceptible to colonization history than another. Yet, knowing this can advance our understanding about the probabilistic nature and predictability of ecological communities.

Here, we provide a theoretical platform upon which to derive general conclusions about the link between colonization history and final community composition. Specifically, we build on the classic Lotka-Volterra (LV) model [27–29] to establish mathematical conditions under which community composition depends on colonization history.

Firstly, to illustrate the scope and assumptions behind our study, we start our analysis by considering a small pool of 3 species that can coexist at a unique equilibrium (see Fig. 1). In this example, there are 6 possible colonization histories (or trajectories) from which an ecological community of 3 species can be assembled by introducing one species at a time, i.e., via successive invasions. If any subset of the species cannot coexist, then the final community composition will surely depend on the colonization history. (Here we assume that the ecological dynamics is fast enough to reach equilibrium before the next species invasion.) For instance, as shown in Fig. 1b, if species 2 and 3 cannot coexist, then the trajectories 3 → 2 → 1 and 2 → 3 → 1 cannot achieve the final community composition {1, 2, 3}, while other trajectories can. Instead, for the system shown in Fig. 1e, any subset of the 3 species can coexist at a unique equilibrium. In this case, the final community composition is independent of the colonization history.

**FIG. 1.**
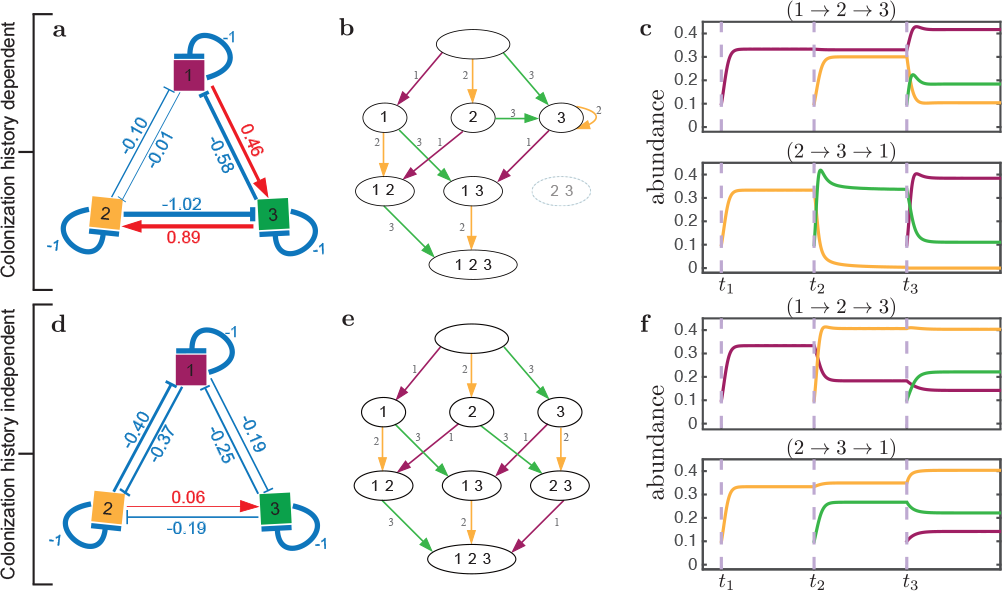
Ecological communities can display different dependencies on colonization history. For illustration purposes, we show the assembly of a 3-species community {1, 2, 3} by the invasion or colonization of one species at a time, following the Lotka-Volterra dynamics. There are in total 3! = 6 different colonization trajectories. (a) The ecological network depicts the pairwise interactions among the three species (which are also encoded in the interaction matrix **A**). The feasible intrinsic growth rate vector **r** is set to be 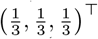. (b) Starting from an empty ecological community (top node), the three species are added successively into the community via different orders. Since species-2 and 3 cannot coexist (gray node), the community composition will be dependent on colonization history. That is, the final state of the three species together cannot be assembled if we follow the trajectory 3 → 2 → 1 or 2 → 3 → 1, while the other four trajectories will lead to the desired final state. (c) As an example, we show two different trajectories and their final community compositions. Panels (d-e) show a similar case as the previous example, but with different interaction matrix **A**. In this case, the community composition is independent on colonization history.

The above observations prompt us to prove two theorems (see Supplemental Material S2 for the proof), which provide sufficient conditions to check whether the final composition of a community of *S* species depends on the colonization history:

#### Theorem 1

If there exists a set of species 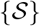 that can coexist at a unique equilibrium, but a smaller subset of species 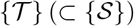 cannot, then the final community composition formed by the 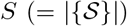 species depends on the colonization history.

#### Theorem 2

If any subsets of species 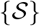 can coexist at a unique equilibrium, then the final community composition formed by the 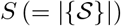 species does not depend on the colonization history.

In this work, we focus on the simple scenario that the *S* species can coexist at a unique equilibrium for two main reasons. First, the *S* species should be able to coexist, otherwise it makes no sense to investigate the impact of colonization history on the final community composition. Second, they should coexist at a unique equilibrium. Otherwise, if the system displays multi-stability, i.e., the equilibrium state depends on initial conditions, then the impact of colonization history on the final community composition becomes trivial. Moreover, if the system undergoes limit cycles or even chaos, then it will be too difficult to investigate the impact of colonization history on the final community composition. Thus, our work provides a first-order approximation to the effect of colonization history on community composition.

To more quantitatively study the impact of colonization history on community composition, we consider ecological systems governed by the classic LV model, which is tractable enough to allow us to formally investigate the conditions under which community composition depends on colonization history. The LV model can be written as follows

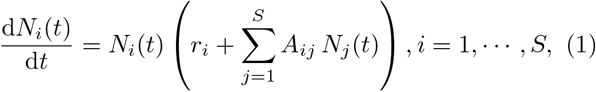

where *N*_*i*_ is the abundance (or biomass) of species-*i*, *S* corresponds to the number of species in the community, A = [A_*ij*_]_*S×S*_ is the interaction matrix whose elements denote the per capita effect of one species on the per capita growth rate of another species, and *r*_*i*_ is the intrinsic growth rate of species-*i*.

To ensure that the *S* species can coexist at a unique equilibrium, following previous studies [30, 31], we focus on diagonally stable interaction matrices **A** (i.e., there is a positive definite diagonal matrix **D** such that **DA** + **A**^⊤^ **D** is a negative definite symmetric matrix [32]). A diagonally stable interaction matrix **A** guarantees that the LV model has a single, globally, attractive equilibrium [33]. We emphasize that the assumption of a diagonally stable interaction matrix **A** is deeply driven by the complexity of this problem and allows us to focus on the feasibility of the system—the necessary condition for species coexistence [34, 35].

To construct a diagonally stable **A** matrix, we randomly choose the off-diagonal elements *A*_*ij*_(*i* ≠ *j*) with probability *C* from a normal distribution with mean zero and standard deviation 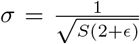. Additionally, the diagonal elements (self-regulation parameters) are set to be *A*_*ii*_ = –*d*. It can be proven that for any *ϵ* > 0 and *d* = 1, the randomly-generated interaction matrix **A** is diagonally stable [36]. Here, we set *ϵ* = 0.01. It is worth noting that *C* represents the connectance of the community (i.e., the ratio between actual and potential interactions in the ecological network), and *σ* denotes the characteristic interspecific interaction strength.

Furthermore, to ensure that the *S* species can coexist at a feasible equilibrium (i.e., all the species abundances are positive) given an interaction matrix **A**, we choose the intrinsic growth rate vector **r** randomly within the feasibility domain *D*_*F*_ (**A**) [37]. The feasibility domain is an algebraic cone described by 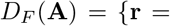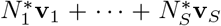 with 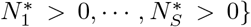 [38]. Note that 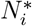 is the equilibrium abundance of species-*i*, and **v**_*i*_ is the spanning vector of the algebraic cone, where the *j*-th component of **v**_*i*_ is given by 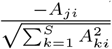 [37]. This procedure guarantees that there is at least one colonization trajectory that can given rise to the whole community formed by *S*-coexisting species. Further information about the parameterization of **A** and **r** can be found in Supplemental Material.

As noted in our two theorems above, whether all subsets of species can coexist at equilibrium will determine if the community composition depends on colonization history. Under the LV dynamics, this coexistence is guaranteed if the equilibria of System (1) are feasible (i.e., all present species have positive abundance) and globally stable for all sub-communities. It has been proved that if the interaction matrix **A** is diagonally stable, then all sub-matrices 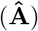 are diagonally stable as well [32], and the non-trivial positive equilibrium point will be globally asymptotically stable (that is, species can stably coexist) [33]. These matrix properties imply that we only need to guarantee the feasibility of the equilibrium for all sub-communities 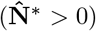. The unique equilibrium of every sub-community under LV dynamics can be calculated as 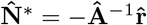, where 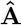 and 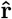 are the subsets of the interaction matrix and the intrinsic growth rate vector of the corresponding sub-community. Thus, to describe the relationship between the final community composition and colonization history, the two theorems can be simplified to the following corollary:

## Corollary 1

For a community 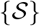 that follows the LV dynamics characterized by a diagonally stable interaction matrix **A** and a feasible intrinsic growth rate vector **r**, if there exists a subset 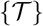 of species that does not have a feasible equilibrium, then the final community composition of the community 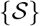 will depend on the colonization history. Otherwise, it will be colonization-history independent.

To illustrate the application of Corollary 1, we consider the two 3-species communities shown in Figure 1. The community shown in Figure 1a is characterized by a feasible intrinsic growth rate vector 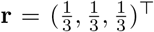, and a diagonally stable interaction matrix

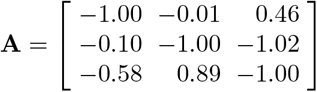

In this case, we can verify that there exists a feasible equilibrium **N**\* = (0.417; 0.104; 0.184)^⊤^ for the three species, but there is no feasible equilibrium for the species-pair {2, 3}. Thus, following Corollary 1, the final community composition will depend on the colonization history. Indeed, Fig. 1c shows two different final states obtained by two colonization trajectories. Figure 1d shows a community characterized by the same feasible intrinsic growth rate vector **r**, but with a different diagonally stable interaction matrix

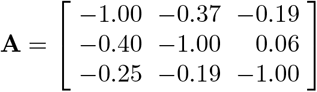

In this case, all subsets of species have a unique and feasible equilibrium. Thus, the final community composition does not depend on the colonization history: any order of species arrival will eventually yield the same final community composition of the three species (see Fig. 1f for examples of two orders).

Next, we perform extensive numerical simulations to investigate which properties of the community and individual species can increase the probability that community composition is dominated by colonization history. Specifically, we study the effect of community properties by systematically generating diagonally stable interaction matrices with different community size (*S*), network connectance (*C*), and interaction types. To investigate the extent to which the variation of intrinsic properties across species affect the link between colonization history and community composition, we systematically generate feasible vectors of intrinsic growth rates **r** with different levels of variability across the elements. In sum, we sample 2, 000 feasible vectors **r** ∊ *D*_*F*_ (**A**) for each randomly-generated interaction matrix **A** across a mixed gradient of community size, connectance, and interaction type. For each point within the gradient, we use an ensemble of 50 different realizations of **A** to calculate an expected value and error bars.

First, we find that the probability that the community composition depends on the colonization history, hereafter denoted as *P*, always increases with the community size *S* (Fig. 2-top) or network connectance *C* (Fig. 2-bottom), regardless of the interaction type. This indicates that the community composition will almost surely depend on the colonization history when an ecological system is composed of a large number of species or when species are highly connected.

**FIG. 2.**
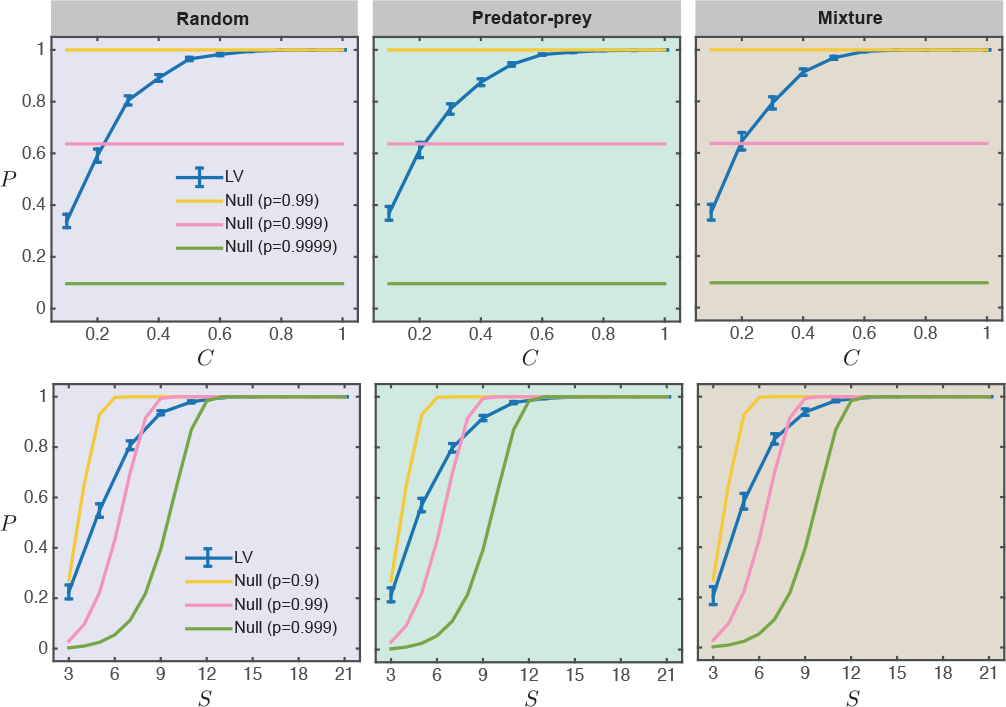
The probability *P* that community composition depends on colonization history as a function of community properties. *P* is calculated as a function of community properties: community size *S*, connectance *C*, and interaction types, using the LV model (line plots with error bars) or the null model (lines). (Top) *P* as a function of connectance *C*, with fixed community size *S* = 10. (Bottom) *P* as a function of community size *S*, with fixed connectance *C* = 0.5. Each column corresponds to a particular interaction type: random interactions, predator-prey, and mixture of competition and mutualism.

To check how this result holds beyond expectations, we consider a simple null model as follows.We assume that a random subset of species has a fixed probability *p* to have a feasible equilibrium. Then, the probability to have at least one subset of species that does not have a feasible equilibrium is given by 1 – *p*^*n*^, where *n* = 2^*S*^ − 2 is the number of possible subsets with at least one and at most (*S* – 1) species. According to Corollary 1, we conclude that the probability that the community composition depends on the colonization history is *P*_null_ = 1 − *p*^*n*^, where the subscript ‘null’ stands for the null model.

In the top panel of Fig. 2, for a given community size *S*, we plot *P*_null_ based on different values of *p* (horizontal lines).Clearly, lower *p* yields higher *P*_null_, regardless of the connectance *C*. For example, for *S* = 10, we have *P*_null_ ~ 0.1 for *p* = 0.9999 (green line), and *P*_null_ ~ 1 for *p* = 0.99 (yellow line). However, our calculation based on the LV model indicates that *P* increases monotonically with increasing *C*, and for *S* = 10 we have *P* → 1 only if *C* is above 0.6, regardless of the interaction types. In the bottom panel of Fig. 2, for a given value of *p*, we plot *P*_null_ as a function of the community size *S*, finding that *P*_null_ increases monotonically with *S*. Note that the *S*-dependency of *P*_null_ is heavily driven by the value of *p*. For example, for *p* = 0.999 (or 0.9), *P*_null_ will always underestimate (or overestimate, respectively) *P* for *S* < 12, regardless of the interaction types. The difference between *P*_null_ and *P* shown here suggests that ecological dynamics (as simple as they can be) can fundamentally alter the dependency of the community composition on colonization history. That is, this dependency cannot be precisely predicted from the probability of feasibility of each individual subset (as assumed in the null model).

Second, we find that the higher the interspecific variation (i.e., the variation of species’ intrinsic growth rates) within a community, the higher the probability that community composition depends on colonization history. We quantify the level of interspecific variation as the variation of intrinsic growth rates within the feasible vector, i.e., 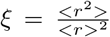 with **r** *∈ D*_*F*_ (**A**). Figure 3 shows that *P* increases monotonically with increasing *ξ*, regardless of the interaction type. In particular, when all species have the same intrinsic growth rate, rendering the lowest level of variation *ξ* = 1, colonization history has the weakest effect on the final community composition. But, as soon as intrinsic growth rates start to differ among species, colonization history becomes a dominant force modulating the final community composition. Here, for the given community size *S*, we also plot *P*_null_ based on different values of *p*. As shown in Fig. 3 (horizontal lines), lower *p* yields higher *P*_null_, regardless of the inter-specific variation *ξ* and interaction types. For example, for *S* = 8, we have *P*_null_ ~ 0.01 for *p* = 0.9999 (green line), and *P*_null_ ~ 0.9 for *p* = 0.99 (yellow line). However, our calculation based on the LV model indicates that *P* increases monotonically with increasing *ξ*, and for *S* = 8 we have *P* → 1 only if *ξ* is above 2, regardless of the interaction types. The difference between *P*_null_ and *P* underscores the impact of interspecific variation on the probability that community composition depends on colonization history, which cannot be predicted from the null model.

**FIG. 3.**
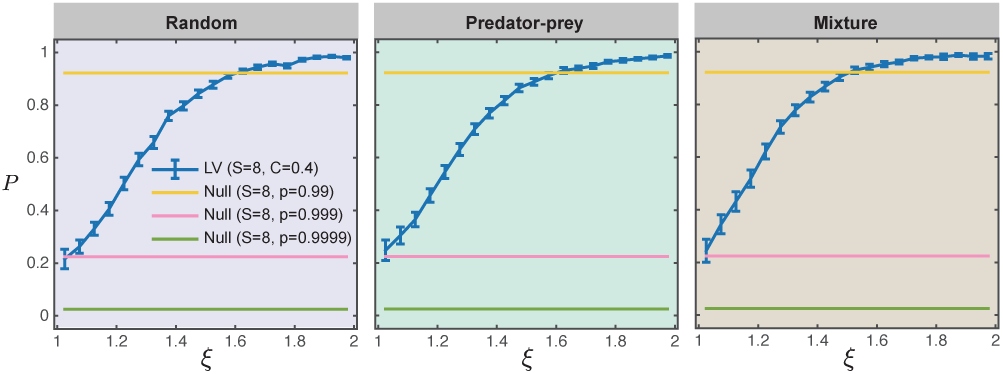
The probability *P* that community composition depends on colonization history as a function of intrinsic properties of species. The probability is calculated for different levels of interspecific variation (intrinsic growth rate) and interaction types, using the LV model (line plots with error bars) or the null model (lines). We fix community size *S* = 8 and network connectance *C* = 0.4. We arbitrarily sampled feasible vectors of intrinsic growth rates and categorized them (bin width=0.05) according to the variation (*ξ*) across their elements: 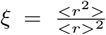. We used the curves to replace the histogram and showed the error bars for 50 different cases. Each column corresponds to a particular interaction type: random interactions, predator-prey, and mixture of competition and mutualism.

In summary, here we have offered a testable theory linking colonization history and community composition. In particular, we have proved two theorems and one corollary to determine whether the composition of a community governed by LV dynamics will depend on its colonization history. Moreover, we have demonstrated that the probability that community composition is dominated by colonization history increases monotonically with community size, network connectance, and variation of intrinsic growth rates across species. These effects cannot be precisely predicted from a simple null model that only considers the probability of feasibility of each each sub-community of species. It is worth noting that our current framework has several limitations. First, it focuses on the coexistence of species at a globally stable equilibrium based on the assumption of a diagonally stable interaction matrix. Note however that global stability has been considered on the most relevant ecological cases [39]. In fact, when the interaction matrix is not diagonally stable, the LV model will be trivially history dependent. That is, if a system depends on the initial value, it can perform multi-stability so that its final composition will always depend on the colonization history [40]. Second, because the coexistence of species not only could be driven by an equilibrium state, but also could be driven by different dynamical properties, such as limit cycles or chaotic behavior, those more complicated scenarios deserve a more dedicated research effort [41]. Third, the simulation framework is applicable to LV dynamics with linear functional responses. Extending the calculations to population dynamics models with more complicated functional response will also be an interesting future direction [35, 42]. Despite those limitations, the simplicity of our work have allowed us to provide a first-order classification of the conditions modulating the impact of colonization history. For example, this work can serve as a basis for future work aiming to study the extent to which it is possible to reconstruct (or to partially reconstruct) the species arrival order in a community. Overall, our work provides a formal probabilistic understanding of the complex process of community assembly.

## References

[1] J. M. Diamond, Assembly of species communities, in Ecology and Evolution of Communities (Harvard University Press, Cambridge, MA, 1975) pp. 342–444.

[2] E. P. Odum, The strategy of ecosystem development, in The Ecological Design and Planning Reader (Springer, 2014) pp. 203–216.

[3] M. Vellend, The theory of ecological communities (MPB-57) (Princeton University Press, 2016).

[4] R. M. May, Science 241, 1441 (1988).

[5] T. Fukami, Community assembly dynamics in space, in Community Ecology: Processes, Models, and Applications (Oxford University Press, Oxford, 2010) pp. 45–54.

[6] T. Fukami, Annu. Rev. Ecol. Evol. Syst. 46, 1 (2015).

[7] D. Tilman, Resource competition and community structure (Princeton university press, 1982).

[8] P. J. Morin, Ecology 65, 1866 (1984).

[9] M. Jaarola, H. Tegelström, and K. Fredga, Biol. J. Linnean Soc. 68, 113 (1999).

[10] D. M. Fonseca and D. D. Hart, Ecology 82, 2897 (2001).

[11] I. Martínez, M. X. Maldonado-Gomez, J. C. Gomes-Neto, H. Kittana, H. Ding, R. Schmaltz, P. Joglekar, R. J. Cardona, N. L. Marsteller, and S. W. Kembel, Elife 7, e36521 (2018).

[12] E. Weiher and P. Keddy, Ecological assembly rules: perspectives, advances, retreats (Cambridge University Press, 2001).

[13] J. M. Chase, Oecologia 136, 489 (2003).

[14] T. Fukami, Ecology 85, 3234 (2004).

[15] D. Sprockett, T. Fukami, and D. A. Relman, Nat. Rev. Gastroenterol. Hepatol. 15, 197 (2018).

[16] J. A. Drake, Am. Nat. 137, 1 (1991).

[17] L. R. Belyea and J. Lancaster Oikos 86, 402 (1999).

[18] J. M. Chase, Proc. Natl Acad. Sci. USA 104, 17430 (2007).

[19] L. Götzenberger, F. de Bello, K. A. Bråthen, J. Davison, A. Dubuis, A. Guisan, J. Lepš, R. Lindborg, M. Moora, M. Pärtel, L. Pellissier, J. Pottier, P. Vittoz, K. Zobel, and M. Zobel, Biol. Rev. 87, 111 (2012).

[20] C. Song, F. Altermatt, I. Pearse, and S. Saavedra, Ecol. Lett. 21, 1221 (2018).

[21] J. E. Price and P. J. Morin, Ecology 85, 1017 (2004).

[22] C. F. Steiner and M. A. Leibold, Ecology 85, 107 (2004).

[23] W. M. Post and S. L. Pimm, Math. Biosci. 64, 169 (1983).

[24] J. A. Drake, J. Theor. Biol 147, 213 (1990).

[25] R. Law and R. D. Morton, Ecology 74, 1347 (1993).

[26] J. L. Lockwood, R. D. Powell, M. P. Nott, and S. L. Pimm, Oikos 80, 549 (1997).

[27] A. J. Lotka, Elements of physical biology (Williams and Wilkins, Baltimore, 1925).

[28] V. Volterra, Nature 118, 558 (1926).

[29] T. Yasuhiro, Global dynamical properties of Lotka-Volterra systems (World Scientific, 1996).

[30] S. Saavedra, R. P. Rohr, J. Bascompte, O. Godoy, N. J. Kraft, and J. M. Levine, Ecol. Monogr. 87, 470 (2017).

[31] C. A. Serván, J. A. Capitán, J. Grilli, K. E. Morrison, and S. Allesina, Nat. Ecol. Evol. 2, 1237 (2018).

[32] E. Kaszkurewicz and A. Bhaya, Matrix diagonal stability in systems and computation (Springer Science and Business Media, 2012).

[33] B. S. Goh, Am. Nat. 111, 135 (1977).

[34] J. Hofbauer and K. Sigmund, Evolutionary games and population dynamics (Cambridge University Press, 1998).

[35] S. Cenci and S. Saavedra, Phys. Rev. E 97, 012401 (2018).

[36] T. E. Gibson, A. Bashan, H.-T. Cao, S. T. Weiss, and Y.-Y. Liu, PLoS Comput. Biol. 12, e1004688 (2016).

[37] C. Song and S. Saavedra, Ecology 99, 743 (2018).

[38] C. Song, R. P. Rohr, and S. Saavedra, J. Theor. Biol. 450, 30 (2018).

[39] R. C. Lewontin, Brookhaven Symp. Biol. 22, 13 (1969).

[40] G. Bunin, Phys. Rev. E 95, 042414 (2017).

[41] S. J. Schreiber, M. Yamamichi, and S. Y. Strauss, Ecology 100, e02664 (2019).

[42] M. AlAdwani and S. Saavedra, Math. Biosci. 315, 108222 (2019).

