## Supplementary Information for "The impact of colonization history on the composition of ecological systems"

### Supplemental Material for: The impact of colonization history on the composition of ecological systems

#### S1 Feasibility and stability of fixed points

The stable coexistence of species means that not only the fixed point (or equilibrium point) is a feasible solution, but also it is globally stable. For this purpose, the fixed point of an LV System (see main text) can be calculated via

$$N_i(t)\left(r_i + \sum_{j=1}^{S} A_{ij} N_j(t)\right) = 0. \tag{1}$$

In general, there are $2^S$ possible roots for Eq. (1), including $N_i = 0$, which denote the theoretical final state of presence and absence of each species. Note that only one of these roots can lead to a feasible fixed point (i.e., $N_i^* > 0$) [1, 2]. Under LV dynamics, if the final community includes $k$ species after an invasion, the theoretical fixed point for the presence of $k$ species can be obtained as follows

$$\mathbf{N}^{(k)} = -\left(\mathbf{A}^{(k)}\right)^{-1} \mathbf{r}^{(k)}. \tag{2}$$

Here $\mathbf{A}^{(k)}$ ($k \times k$ matrix) and $\mathbf{r}^{(k)}$ ($k$-dimensional vector) are the corresponding interaction matrix and growth rate of the existing k species respectively. Note that $\mathbf{A}^{(k)}$ needs to be invertible. To determine feasibility, the fixed point $\mathbf{N}^{(k)}$ needs to satisfy $\mathbf{N}^{(k)} > \mathbf{0}$ (i.e., every component is positive). For the stability of the fixed point, we assume that the matrix $\mathbf{A}$ is diagonally stable (constructed by the method in Ref. [3], where we can confirm that $\mathbf{A}$ is also invertible), and its principal submatrices $\mathbf{A}^{(k)}$ are also diagonally stable [4]. Hence, the feasible solutions obtained by Eq. (2) are always globally stable [5].

#### S2 Proof of Theorems 1 and 2

**Theorem 1**: If there exists a set of species $\{\mathcal{S}\}$ that can coexist at a unique equilibrium, but a smaller subset of species $\{\mathcal{T}\}$ ($\{\mathcal{T}\} \subset \{\mathcal{S}\}$) cannot, then the final community composition formed by the $S$ ($S = |\{\mathcal{S}\}|$) species depends on the colonization history.
**Proof:**. Let $|\{\mathcal{S}\}| = S$, and $|\{\mathcal{T}\}| = T$. Without loss of generality, let us consider $\{\mathcal{T}\} = \{1, 2, ..., T\}$, and $\{\mathcal{S}\} = \{1, ..., T, T+1, ..., S\}$. Since the first $T$ species cannot coexist, if we colonize them first, some of them will become extinct. If we then colonize species $T+1, ..., S$, then the final state will be quite different from the case when we introduce all the $S$ species together.
**Theorem 2**: If any subsets of species $\{\mathcal{S}\}$ can coexist at a unique equilibrium, then the final community composition formed by the $S$ ($S = |\{\mathcal{S}\}|$) species does not depend on the colonization history.
**Proof:**. Without loss of generality, when new species are added in a pool to build a new community $\{\mathcal{I}\} = \{1, 2, ..., i\}$, the community with $i$ species can stably coexist in the unique final state $\hat{\mathbf{N}}^* = (\hat{N}_1^*, \hat{N}_2^*, ..., \hat{N}_i^*)$, which is stable and feasible ($\hat{N}_j^* > 0, j = 1, 2, \cdots, i$). Following these steps to assemble more species, we will find that every final state in the corresponding subsystems is always fixed and unique. Thus, the final state is not dependent on the colonization history.

#### S3 Generating interaction matrices A

All matrices were generated by first building a binary structure as Erdős-Rényi (ER) random networks [6]. For this purpose, first we begin with $S$ isolated nodes (species), we select a node pair, and produce a random number between 0 and 1. If the random number is greater than $C$ (here $C$ is called connectance of the network), we use an edge to connect the above node pair, otherwise the pair remains disconnected. We repeat the above step for each of the $\frac{[S(S-1)]}{2}$ node pairs. Once networks were constructed, we establish the interaction types (signs) between each pair of species. We consider three interaction types: random interaction (no sign structure), predator-prey $(+,-)$, and mixture of competition $(-,-)$ and mutualism $(+,+)$. As mentioned in the main text, to ensure the diagonal stability of the interaction matrix $\mathbf{A}$, we drew the off-diagonal elements $A_{ij}(i \neq j)$ from a normal distribution $\mathcal{N}\left(0, \frac{1}{S(2+\epsilon)}\right)$, and the diagonal elements $A_{ii}$ are set to $A_{ii} = -d$. Additionally, we perform the following procedure to ensure the interaction types. To establish random interactions, if there is an edge between node $i$ and node $j$ $(i, j = 1, 2, \cdots, S)$ (i.e., $A_{ij} = 1$), $A_{ij}$ are selected independently from the normal distribution $\mathcal{N}\left(0, \frac{1}{S(2+\epsilon)}\right)$, and all diagonal entries remain unchanged ($A_{ii} = -d$). To establish predator-prey interactions, if $A_{ij} = 1(i > j)$, we need to find a random number $p$ from a uniform distribution $U[0, 1]$ . If $p \leq 0.5$, $A_{ij}$ is drawn from a half-normal distribution $\left|\mathcal{N}\left(0, \frac{1}{S(2+\epsilon)}\right)\right|$, and $A_{ji}$ belongs to a negative half-normal distribution $-\left|\mathcal{N}\left(0, \frac{1}{S(2+\epsilon)}\right)\right|$. If $p > 0.5$, we do the opposite criterion. To establish a mixture of competition and mutualism, if $A_{ij} = 1(i > j)$, we choose a random number $p$ from a uniform distribution $U[0, 1]$. If $p \leq 0.5$, $A_{ij}$ and $A_{ji}$ are drawn independently from a half-normal distribution$\left|\mathcal{N}\left(0, \frac{1}{S(2+\epsilon)}\right)\right|$. If $p > 0.5$, $A_{ij}$ and $A_{ji}$ are drawn from a negative half-normal distribution$-\left|\mathcal{N}\left(0, \frac{1}{S(2+\epsilon)}\right)\right|$.

### S4 Generating feasible vectors of intrinsic growth rates

To ensure the coexistence of the whole community with $S$ species, we use the feasibility domain to construct feasible vectors of intrinsic growth rates [7]. The feasibility domain can be determined totally when the interaction matrix $\mathbf{A}$ is given. That is, it can be described by

$$D_F(\mathbf{A}) = \{\mathbf{r} = N_1^*\mathbf{v}_1 + \cdots + N_S^*\mathbf{v}_S, \text{with} \quad N_1^* > 0, \cdots, N_S^* > 0\}, \tag{3}$$

where $N_i^*$ is the equilibrium abundance of species $i$, and $\mathbf{v}_i$ is the spanning vector, whose $jth$ component is the normalized column vector $jth$ of the interaction matrix $\mathbf{A}$, i.e., $v_{ij} = \frac{-A_{ji}}{\sqrt{\sum_{k=1}^{S} A_{ki}^2}}$. Thus, if the vector of intrinsic growth rates $\mathbf{r}_f$ is chosen inside the feasibility domain $D_F(\mathbf{A})$, the community with $S$ species will always be feasible. This feasible vector can be defined as follows:

$$\mathbf{r}_f = \sum_{i=1}^{S} N_i^*\mathbf{v}_i, \tag{4}$$

where $N_i^*(i = 1, ..., S)$ are all values in (0,1), and $\sum_{i=1}^{S} N_i^* = 1$.

### S5 Calculating the probability that community composition is dependent on colonization history

Given a set of conditions assumed for an interaction matrix $\mathbf{A}$, we randomly generated 50 different realizations of matrix $\mathbf{A}$. For each realization, we used its feasible domain $D_F(\mathbf{A})$ to extract 100 feasible vectors of intrinsic growth rates $\mathbf{r}$ following the LV model (1). Then, we determined the relationship between final community composition and colonization history via Corollary 1. That is, for each matrix, we calculated the relative frequency (using the feasible vectors) that the final state depends on colonization history. Then, we used the ensemble of 50 different realizations of $\mathbf{A}$ to calculate a mean and error bars for this probability.

#### References


[1] Takeuchi Yasuhiro. *Global dynamical properties of Lotka-Volterra systems.* World Scientific, 1996.

[2] Mohammad AlAdwani and Serguei Saavedra. Is the addition of higher-order interactions in ecological models increasing our understanding of ecological dynamics? *Math. Biosci.*, 35:108222, 2019.

[3] Travis E Gibson, Amir Bashan, Hong-Tai Cao, Scott T Weiss, and Yang-Yu Liu. On the origins and control of community types in the human microbiome. *PLoS Comput. Biol.*, 12(2):e1004688, 2016.

[4] Eugenius Kaszkurewicz and Amit Bhaya. *Matrix diagonal stability in systems and computation.* Springer Science and Business Media, 2012.

[5] Bo S Goh. Global stability in many-species systems. *Am. Nat.*, 111(977):135–143, 1977.

[6] Paul Erdős and Alfréd Rényi. On the evolution of random graphs. *Publ. Math. Inst. Hung. Acad. Sci*, 5(1):17–60, 1960.

[7] Chuliang Song and Serguei Saavedra. Will a small randomly assembled community be feasible and stable? *Ecology*, 99(3):743–751, 2018.
